# In situ Fucosylation for Modulating Wnt Signaling in Live Cells

**DOI:** 10.1101/726612

**Authors:** Senlian Hong, Lei Feng, Hao Jiang, Xiaomeng Hou, Peng Guo, Florence L. Marlow, Pamela Stanley, Peng Wu

## Abstract

Wnt/β-catenin signaling, also known as canonical Wnt signaling, regulates critical, context-dependent transcription in numerous (patho) physiological events. Amongst the well-documented mechanisms of canonical Wnt signaling, modification of N-glycans by L-fucose (Fuc) is the newest and the least understood. Using a combination of Chinese Hamster Ovary (CHO) cell mutants with different fucosylation levels and *in situ* cell-surface Fuc editing (ISF), we report that α(1-3)-fucosylation of N-acetylglucosamine in the LacNAc (Galβ(1-4)-GlcNAc) sequences of complex N-glycans modulates Wnt signaling by regulating the endocytosis of low density lipoprotein receptor-related protein 6 (LRP6). Pulse-chase experiments reveal that increasing N-glycan LacNAc fucosylation elevates endocytosis of lipid-raft-localized LRP6, leading to the suppression of Wnt-β-catenin signaling. Inhibiting endocytosis by inhibiting dynamin 1, a GTPase responsible for endocytosis in eukaryotic cells, partially rescues Wnt signaling. Remarkably, inhibition of Wnt signaling by N-glycan LacNAc fucosylation is fully rescued by the addition of free Fuc to the medium, suggesting that endocytosis of N-glycan fucosylated LRP6 may be mediated by a receptor that recognizes the bound α(1-3)-Fuc. This work provides the first evidence that *in situ* cell-surface fucosylation can be exploited to regulate a specific signaling pathway via endocytosis, revealing a novel regulatory mechanism linking glycosylation of a cell surface receptor with its intracellular signaling.

## Introduction

In the modern era of molecular medicine, prodigious effort has been devoted to dissecting the functions of signaling pathways and the underlying molecular mechanisms. Despite the known involvement of the glycans on cell-surface glycoproteins in numerous signaling pathways, their specific contributions to pathway activities are understood in only a few cases, including O-glycans regulation of Notch signaling^1^ and sialic acid and Fuc regulation of EGF receptor signaling^2^.

It is challenging to study the impact of glycosylation on a specific signaling pathway. This is partly because glycosylation is a posttranslational modification and not under direct genetic control. Most methods to manipulate glycosylation patterns alter glycosylation status globally rather than on a particular or a restricted number of glycoproteins. However, by overexpressing a protein of interest, or mutating its glycosylation sites, the impact of glycans on protein functions in a specific pathway can be singled out. Likewise, using a combination of glycan engineering tools and inhibitors of a particular signaling pathway, the specific roles of glycans in modulating this pathway can be elucidated by monitoring signaling output.

Wnt signaling pathway is an evolutionarily conserved pathway that regulates cell fate determination, cell migration, cell polarity, neural patterning and organogenesis during embryonic development.^3,4^ Aberrant regulation of this pathway is associated with a variety of diseases, including cancer, fibrosis and neurodegeneration.^3,4^ The Wnt pathway is commonly divided into β-catenin dependent (canonical) and independent (non-canonical) signaling.^3,4^ The canonical pathway is activated upon binding of secreted Wnt ligands (such as WNT3a and WNT1) to Frizzled receptors (FZD) and co-receptors LRP5 (low-density lipoprotein receptor-related protein 5) or LRP6.^5–7^ The binding relays a signal through Dishevelled, resulting in stabilization and nuclear entry of β-catenin. In the nucleus, β-catenin forms an active complex with LEF (lymphoid enhancer factor) and TCF (T-cell factor) families of transcription factors to promote specific gene expression.

Previously, we discovered that increasing cell surface N-glycan fucosides through the over-expression of the GDP-Fuc transporter SLC35C1 (solute carrier 35C1, a GDP-Fuc transporter) inhibits canonical Wnt signaling during zebrafish embryogenesis.^8^ However, the mechanism by which inhibition occurs remained obscure. Both WNT and WNT receptors are fucosylated,^9,10^ and in certain cancer cell lines such as the lung cancer cell line A549, LRP6 is also fucosylated (Fig. S13). Structural analysis revealed that *N*-glycans in both WNT8 and FZD 8 cysteine-rich domains are solvent-exposed and do not appear to contribute directly to WNT-FZD interactions.^9^ Therefore, we sought to determine if changes in cell-surface fucosylation status modulates Wnt signaling by influencing the function of LRP receptors.^8^

Here, using a combination of CHO glycosylation mutants and fucosyltransferase-mediated cell-surface chemoenzymatic *in situ* fucosylation (ISF), we find that increasing cell-surface N-glycan fucosylation enhances the endocytosis of the lipid-raft localized LRP6, which in turn suppresses Wnt/β-catenin signaling. This inhibition can be fully rescued by externally added free fucose, suggesting the existence of a fucose-binding protein that regulates the endocytosis of LRP6. Aberrant glycosylation is a hallmark of cancer, among which changes in fucosylated N-glycans are often observed. Cancer cells may exploit this unique mechanism to enhance Wnt signaling and thus promote the survival and proliferation of tumors.

## Results

### ISF inhibits the canonical Wnt signaling in zebrafish embryos

Previously, we reported that over-expression of the GDP-Fuc transporter SLC35C1 specifically inhibits canonical Wnt signaling by increasing the levels of cell-surface N-linked fucosides.^8^ However, manipulating the SLC35C1 level is an indirect means of regulating fucosylation of cell-surface N-glycans. To tune the cell-surface glycan fucosylation level directly, we used a technique we termed *in situ* Fuc editing (ISF). This technique exploits a recombinant *H. pylori* α1-3-fucosyltransferase (FucT) to transfer a Fuc residue from the GDP-Fuc donor to the C3-OH of GlcNAc in a LacNAc unit.^11^ LacNAc is abundantly expressed in complex and hybrid N-glycans. Using this technique, Fuc can be added directly to cell surface glycans of cultured cells and living organisms.^12,13^ However, the formation of the zebrafish enveloping layer (EVL)—the monolayer of cells surrounding zebrafish embryos—prevents the use of ISF to modify inner cells of the embryo covered by the EVL.^14^

The EVL first appears at the blastula stage [4.66 hours post fertilization (hpf)] and eventually forms a sealed layer at approximately 90% epiboly (stage 9 hpf).^14^ To modify the inner part of zebrafish embryos, we devised a new strategy to apply ISF at the 8-cell stage (1.25 hpf) of zebrafish embryogenesis by injecting FucT and GDP-Fuc (or its alkyne-tagged analogue GDP-FucAl) directly beneath the chorion (Fig. S1A).^12^ Via this strategy, the ISF system could gain access to all the cells of the embryo and the sealed chorion served as a natural reaction vessel. After 28 °C-incubation for several hours, we fixed the embryos at 7 hpf (70% epiboly stage), and performed the ligand (BTTPS)-assisted copper-catalyzed azide-alkyne [3+2] cycloaddition reaction (CuAAC)^15^ to conjugate a fluorescent probe to FucAl. All cells throughout the treated embryos exhibited strong FITC fluorescence, while no signal was observed in control embryos treated with FucT and GDP-Fuc that could not be labeled (Fig. 1C, and Fig. S1B).

**Figure 1.**
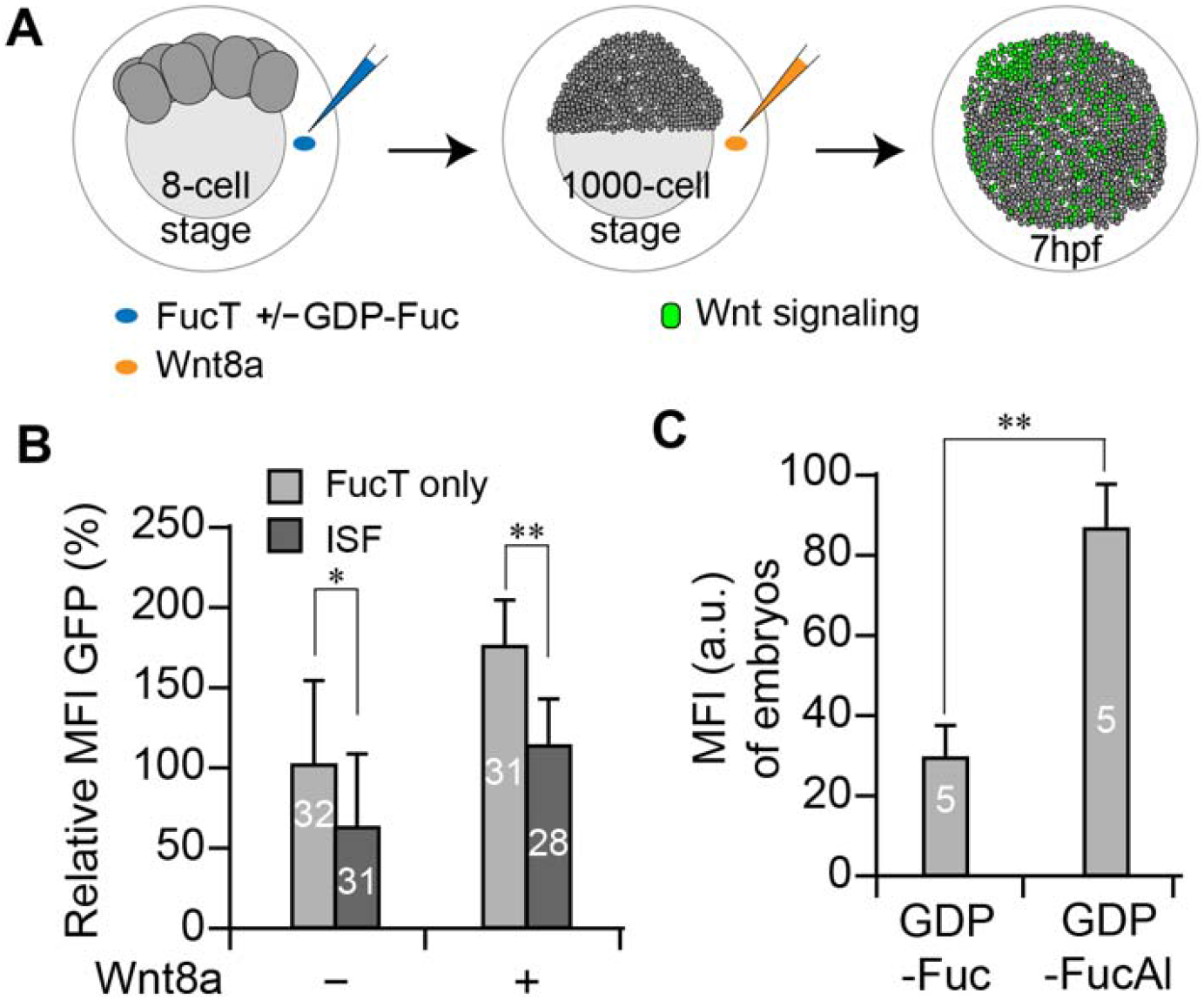
Enhanced α(1-3)-fucosylation via ISF inhibits Wnt signaling in zebrafish embryos. (A) Flow chart of quantifying the impacts of ISF on Wnt signaling activity in a zebrafish Wnt reporter line. At the 8-cell stage, a FucT/GDP-Fuc pre-mixture or FucT only were injected into the chorion to perform ISF or act as control, respectively. To further boost the Wnt signaling readout, recombinant WNT8a was injected into the chorion at the 1000-cell stage before the embryos were transferred individually into single wells. Expression of the GFP reporter was quantified at 7 hpf. (B) Comparing the GFP expression level of embryos treated by ISF or FucT only in the presence or absence of Wnt8a. The relative mean fluorescence intensity (MFI) of GFP was quantified for healthy live embryos at 7 hpf by subtracting background signals with no GFP expression samples, and the GFP-MFI of embryos treated with FucT only was set as 100%. The numbers on the bar graph represent the numbers of healthy embryos used in this assay. (C) quantifying MFI of traditional bioorthogonal chemical reporter labeling in the FucT/GDP-Fuc or FucT/GDP-FucAl treated embryos by ImageJ. Note: * Student’s t test p<0.05; **, Student’s t test p<0.005.

Signaling reporter lines have been used extensively in zebrafish research.^16,17^ Currently, the most sensitive, stable transgenic zebrafish line carrying a Tcf/Lef-miniP:dGFP Wnt signaling reporter was generated by the Ishitani group.^18^ In this reporter line, GFP is detectable in all known Wnt/β-catenin signaling-active cells during embryogenesis starting as early as 3.7 hpf.^18^ We utilized this reporter line to examine the impact of ISF on Wnt activity by quantifying GFP fluorescence. We first treated embryos with FucT and GDP-Fuc or with FucT only at the 8-cell stage (Fig. 1A). At the 1000-cell stage, or approximately 3 hpf, ISF-modified or unmodified embryos were injected into the chorion with recombinant Wnt8a (500 pg)—the main canonical Wnt ligand naturally expressed during gastrulation^19–22^—or left untreated. After the treatment, each embryo was transferred into a well of a 96-well plate and kept at 28 °C to continue development. GFP-associated fluorescence was quantified at 7 hpf. As shown in Fig. 1B, in the presence of exogenously added Wnt8a, we observed stronger Wnt-signaling associated GFP fluorescence. However, embryos modified by ISF exhibited decreased GFP fluorescence compared with control embryos treated with FucT only.

### N-glycan LacNAc fucosylation level is negatively correlated with the canonical Wnt activity

To elucidate potential mechanisms by which N-fucosylation regulates canonical Wnt signaling, we used a combination of ISF and CHO glycosylation mutants with distinct fucosylation statuses. Four CHO lines, Pro-5 parent^23^, and Lec2^24–26^, Lec8^26–29^ and LEC30^30,31^ glycosylation mutants, were chosen for this study. Typical N-glycan structures on each cell line are depicted in Fig. 2A.^32^ Pro-5 CHO cells are the parental cells from which Lec2, Lec8 and LEC30 were derived.^9,10,33^ Pro-5 cells express Fuc in the N-glycan core and peripheral sialic acid in N-glycan antennae, but no fucosylation is found in terminal or internal LacNAc units of complex N-glycans. Lec2 cells express similar N-glycans to Pro-5 cells, but they terminate mainly in Gal and have very few sialic acid residues due to an inactive CMP-sialic acid transporter. Lec8 cells have an inactive UDP-Gal transporter, and therefore their complex N-glycans terminate in GlcNAc with few, if any, having terminal LacNAc.^32,33,34^ The activation of *Fut4* and *Fut9* genes produces the gain-of-function mutant LEC30, which exhibits a significantly increased level of α1-3-fucosylation of LacNAc in N-glycan antennae.^32^

**Figure 2.**
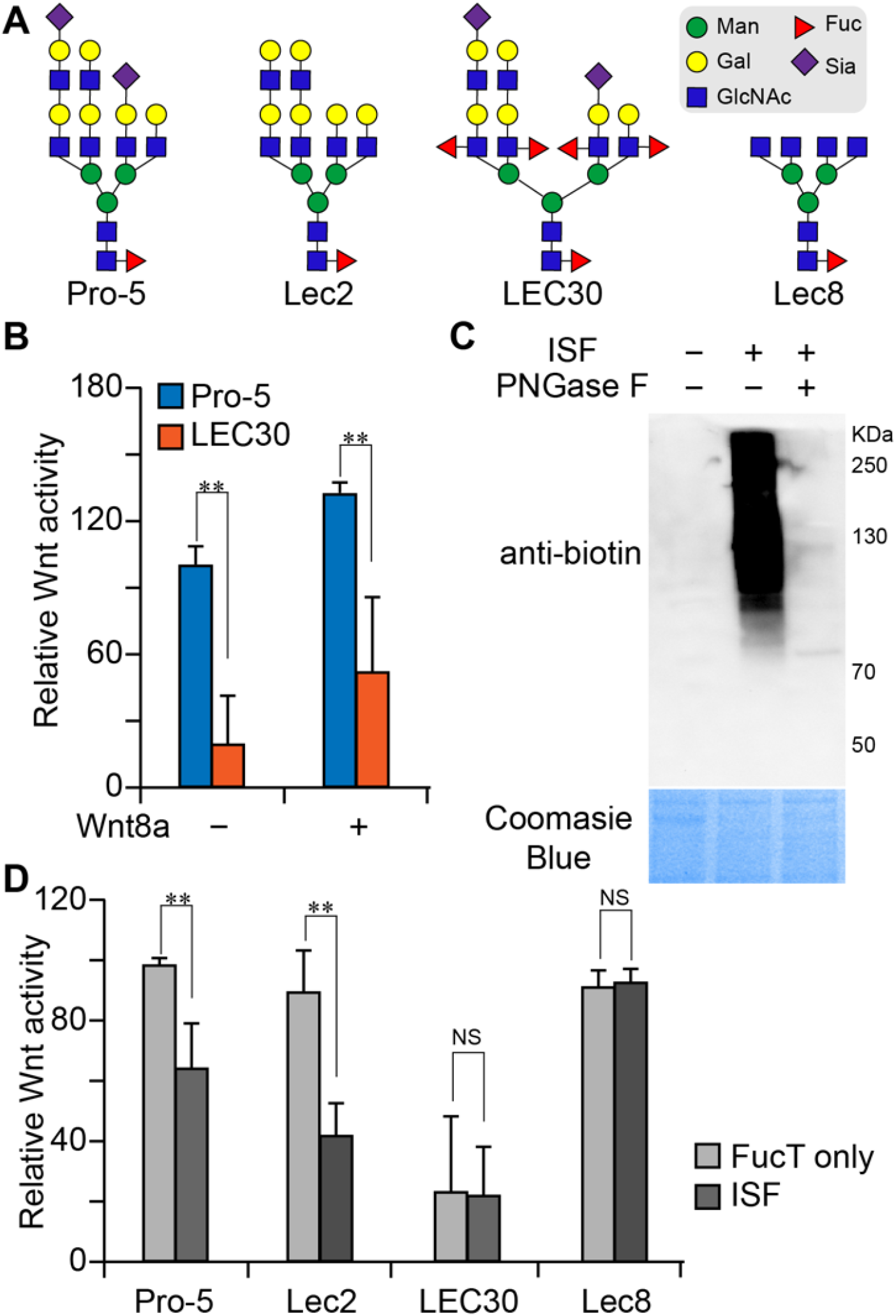
Increased N-glycan LacNAc α(1-3)-fucosylation inhibits Wnt signaling in CHO parent and Lec2 cells. (A) Typical complex N-glycans present on cell surface glycoproteins of CHO Pro-5 cells, and glycosylation mutants Lec2, LEC30 and Lec8 cells.^32^ Pro-5 and Lec2 cells carry LacNAc with no Fuc attached and are excellent substrates for ISF. Lec8 has no LacNAc and LEC30 has many fucosylated LacNAcs. Symbols are from the Symbol Nomenclature for Glycans (SNFG). (B) Comparison of Wnt activity in Pro-5 cells vs. that of LEC30 cells in the presence or absence of Wnt8a. (C) Western blot assay for detection of biotinylated glycoproteins in Pro-5 cells generated via ISF before or after PNGase F-treatment. The equal loading C, was indicted by Coomassie blue staining (lower panel). (D) Comparison of Wnt activity in four CHO parent and three mutant lines treated with ISF or FucT only. Note: **, Student’s t test p<0.005; NS, not significant. The error bars represent the standard deviation of three biological replicates.

To assess if changes in cell-surface N-glycan LacNAc fucosylation have any impact on canonical Wnt signaling in these CHO glycosylation mutants, we examined the effect of ISF on Wnt activity in Pro-5, Lec2 and LEC30 cells. Lec8 cells with no N-glycan LacNAc additions were used as a negative control. The Wnt signaling activity in these cell lines was measured using a luciferase reporter assay,^35^ and the cell-surface α1-3-fucosylation level was characterized using a Fuc-specific lectin, *Aleuria aurantia lectin* (AAL). As shown in Fig. 2, Wnt activity negatively correlated with cell-surface N-glycan LacNAc fucosylation level. In the presence or absence of exogenously added Wnt8a, endogenously fucosylated LEC30 cells exhibited significantly lower Wnt activity compared to Pro-5 cells with no endogenous LacNAc fucosylation (Fig. 2B). After ISF, the cell-surface N-glycans LacNAc fucosylation level in Pro-5 cells was significantly elevated, but the same treatment had almost no effect on the N-glycan LacNAc fucosylation status of LEC30 cells (Fig. 2C, and Fig. S2). Consistent with this observation, Wnt signaling in Pro-5 cells was significantly suppressed by ISF, but the same treatment had little influence on Wnt activity in LEC30 cells (Fig. 2D, and Fig. S3). As expected, ISF had similar impacts on Wnt signaling in both Pro-5 and Lec2 cells. Not surprisingly, no influence on Wnt activity was observed in Lec8 cells following ISF. Finally, using GDP-fucose-biotin as the donor substrate for ISF and PNGase-F to remove N-linked glycans, we confirmed that the newly-added Fuc residues are mainly located in N-glycans of Lec2 and Pro-5 cells (Fig. 2C, and Fig. S4). This was to be expected because these CHO cell lines do not synthesize O-GalNAc glycans with LacNAc.^32^

The above results suggested that Wnt activity is negatively correlated with cell-surface N-glycan LacNAc fucosylation level (Fig. 2D). To further investigate, we performed a time-lapse experiment to evaluate the impact of ISF on Wnt activity. In this experiment, Pro-5 CHO cells were incubated with FucT and GDP-Fuc to perform cell surface ISF. At 30 min intervals, cells were lysed and cell surface fucosylation levels and intracellular β-catenin levels were quantified using Western blot analysis. Alternatively, at each time point, the treated cells were analyzed by the luciferase reporter assay to evaluate Wnt activity. As revealed by quantitative Western blotting analysis, the N-glycan LacNAc fucosylation level increased in the first hour of the treatment, followed by a decline (Fig. 3A) presumably due to the loss of FucT activity after 60 min. As expected, the longer ISF treatment was, the less Wnt activity was ob- served (Fig. 3B), which was accompanied by the continuing decrease of β-catenin levels (Fig. 3C, and 3D).

**Figure 3.**
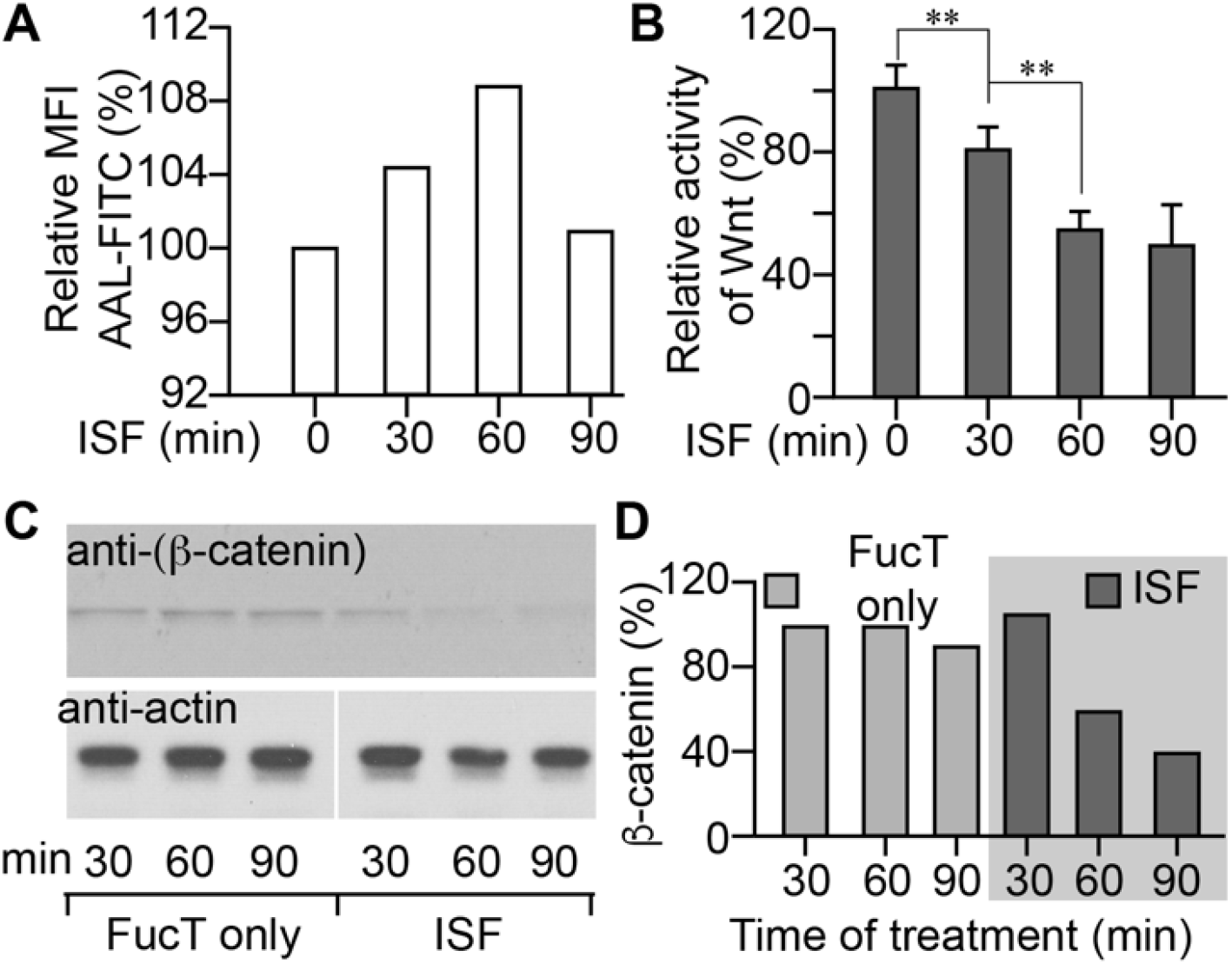
Cell-surface ISF suppresses Wnt/β-catenin signaling in a time dependent manner. (A) Quantitative lectin blot analysis of time-dependent incorporation of α(1-3)-linked Fuc onto Pro-5 cell-surface glycans via ISF-treatment. (B) The time-dependent decrease of Wnt activity in Pro-5 cells treated by ISF. The error bars represent the standard deviation of three biological replicates. In A and B, results are from the same set of samples and the signal intensity of sample at 0 min treatment is set as 100%. (C) Time-dependent decrease of the β-catenin level in ISF-treated Pro-5 cells. Western blot of anti-β-catenin (upper panel) and actin (lower panel). (D) The quantification of anti-β-catenin level in 3C by ImageJ. The 30 min non-treated sample was set as 100%. Note: **, Student’s t test p<0.005.

Next, we blocked the degradation of β-catenin by adding a non-degradable β-catenin or a small-molecule inhibitor, CHIR99021, that inhibits the β-catenin degradation.^36^ Both approaches were effective at enhancing Wnt activity in a dose-dependent manner (Fig. S5). However, neither of these treatments could fully rescue the ISF-induced Wnt inhibition, suggesting that Wnt inhibition caused by ISF is upstream of β-catenin degradation.

### N-glycan LacNAc fucosylation increases LRP6 expression at the cell surface with concomitant Wnt inhibition

Wnt signaling is regulated at multiple levels, but the major mechanism known to degrade Wnt ligand is mediated by LRP6-WNT endocytosis. Tuning the stability and the localization of LRP6 controls Wnt activity.^7^ In general, Wnt activity is positively correlated with cellular LRP6 levels.^6,37,38^ However, in our previous work we found that when Wnt activity was inhibited by elevated N-glycan LacNAc fucosylation induced by SLC35C1 over-expression, the LRP6 protein level was surprisingly increased.

The same unexpected phenomenon was also observed in CHO cells. When Pro-5 cells were subjected to ISF, the total LRP6 protein level was elevated along with the increased cell surface N-glycan LacNAc fucosylation (Fig. 4B). This observation suggested that LRP6 expression or stability at the cell surface may be linked to N-glycan LacNAc fucosylation.

**Figure 4.**
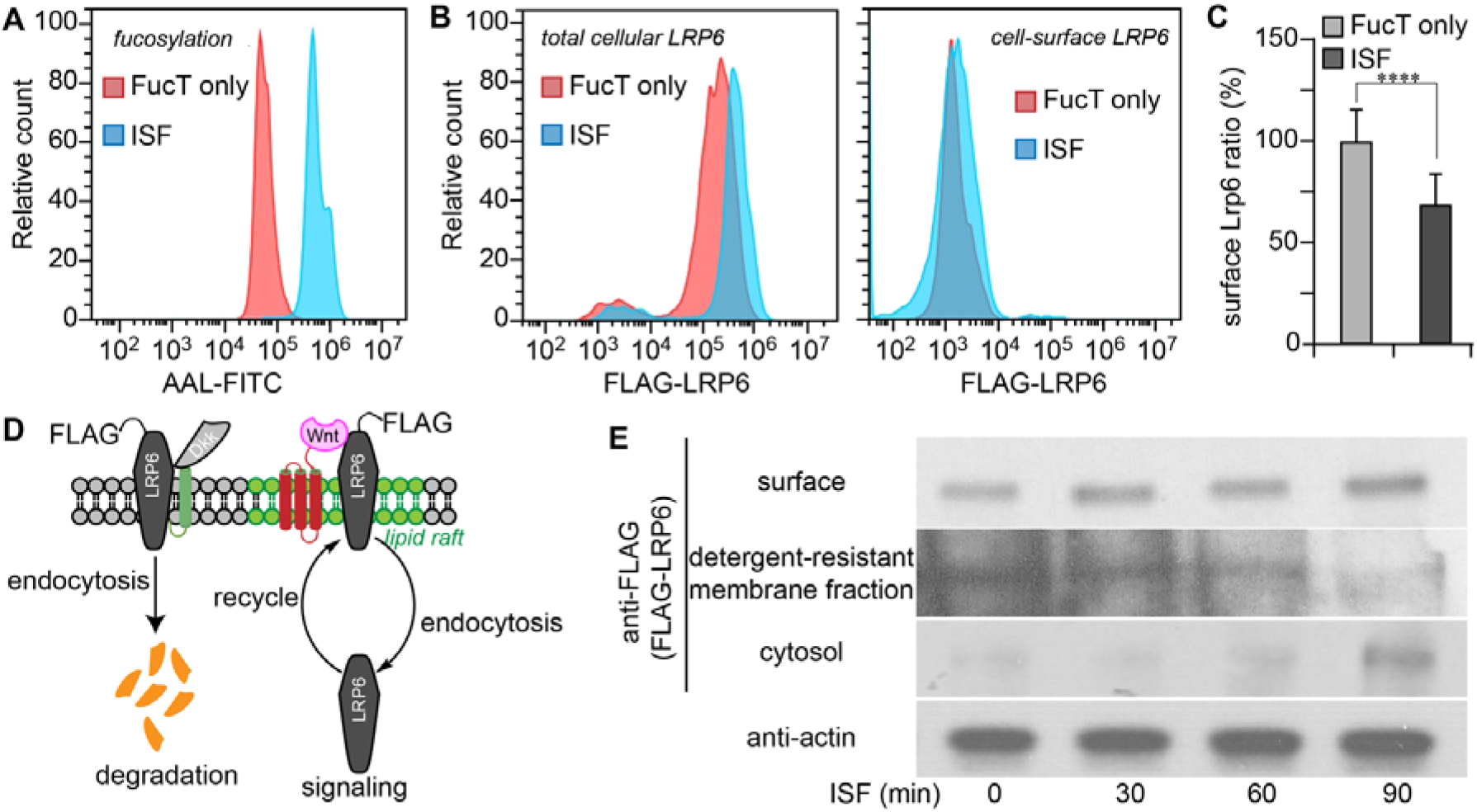
ISF treatment relocates LRP6 from detergent-resistant membranes to cytoplasm (A, B) Flow cytometry histograms of the FucT-treated (Red) and ISF-treated (Blue) Pro-5 CHO cells. The LRP6 in 4B is transiently-expressed FLAG-LRP6. (A) Detection of cell surface N-glycan LacNAc fucosylation by the lectin AAL-FITC. (B) The immunofluorescent staining of cellular overall LRP6 and cell-surface LRP6 level. (C) The ratio of LRP6 (surface to total) in Pro-5 cells treated by ISF, or FucT only. The mock control was set as 100%. Error bars show standard error. N=53,664 cells and p<0.0001. (D) Scheme of LRP6 location and maintenance is regulated through endocytosis in cells. (E) Western blot assay of membrane LRP6, lipid raft LRP6, cytosol LRP6 and actin after different time of ISF treatments.

### N-glycan LacNAc fucosylation relocates the LRP6 protein from detergent-resistant membrane to the cytoplasma

As presently understood, LRP6 stability is controlled by Dickkopf (Dkk) through a specific endocytosis pathway. Dkk antagonizes Wnt signaling by binding to Kremen and LRP6, forming a complex, which is then endocytosed through the clathrin-mediated endocytosis, leading to LRP6 degradation.^38^ On the other hand, the complex formed by Wnt with LRP6 and Frizzled is endocytosed through the caveolin-mediated endocytosis pathway.^5,39^ After sending a signal to inhibit β-catenin degradation, LRP6 is recycled (Fig. 4D).^7^ We hypothesized that N-glycan LacNAc fucosylation may elevate the LRP6 protein level by affecting endocytosis or degradation.

To test these hypotheses, we first assessed the cellular distribution of LRP6 in Pro-5 CHO cells before and after ISF using both flow cytometry and Western blot analyses. For flow cytometry analysis, a N-terminal FLAG-tagged LRP6 was used.^40^ The antibody staining of FLAG on live cells detected surface-localized LRP6, while detection of total LRP6 was determined following fixation and permeabilization of the cells. Raft proteins in the detergent-resistant fraction^41,42^ were re-suspended in 10% SDS buffer and isolated using streptavidin-beads immunoprecipitation. fixed and permeabilized cells revealed overall LRP6 level. With cell surface N-glycan LacNAc fucosylation increased by ISF (Fig. 4A), the total amount of cellular LRP6 was doubled (Fig. 4B). At the same time, cell-surface LRP6 increased only slightly and this increase was not significant. This unexpected result indicated that the increased LRP6 was primarily located inside the cell. Determining the ratio of surface to total LRP6 after ISF revealed that the relative proportion of cell surface LRP6 had decreased by approximately 30% (Fig. 4C).

It was reported that only the LRP6 residing in the lipid rafts can forms a complex with WNT and FZD.^7,43^ We examined the distribution of LRP6 on the cell surface, in lipid rafts and inside the cell using differential biotinylation and Western blot analysis. At various time points following ISF, cell-surface protein was biotinylated. After cell lysis, RIPA buffer was used to extract the soluble fraction, separating the insoluble, lipid raft fraction. We then used streptavidin-beads to enrich biotinylated proteins from the soluble fraction. Finally, lipid raft proteins in the insoluble fraction were resuspended in 10% SDS buffer and isolated using anti-biotin immunoprecipitation.

Western blot analysis revealed that the cell surface LRP6 increased slightly after ISF (Fig. 4E top panel). By contrast, the level of LRP6 in the cytosolic fraction increased notably (Fig. 4E third panel), with a concomitant decrease in the detergent-resistant fraction (i.e., lipid-rafts) localized of LRP6. By 1.5 hr following ISF, LRP6 in the detergent-resistant fraction almost disappeared. Together these results and former reports^7^ provided strong evidence that ISF led to the internalization of LRP6 from lipid rafts, presumably via an increase of caveolin-mediated endocytosis.

To further investigate the cellular redistribution of LRP6 induced by ISF, we performed an imaging experiment to analyze the colocalization of LRP6 and Cholera toxin subunit B (CTB),^44^ a marker for lipid rafts (Fig. 5A). The colocalization of LRP6 and CTB was found both on the cell surface and in the cytoplasm. After treating cells with ISF for 30 min, the percentage of cell surface colocalized LRP6 and CTB dropped from approximately 50% per cell in non-treated cells to ~14% per cell in ISF-treated cells (Fig. 5B). Moreover, the overall colocalization also decreased from ~8% per cell to ~2% per cell after ISF (Fig. 5C). Consistent with this observation, the colocalization of LRP6 and CTB was significantly lower in the highly N-fucosylated LEC30 cells compared to Pro-5 cells that have no N-glycan LacNAc fucosylation (Fig. 5D).

**Figure 5.**
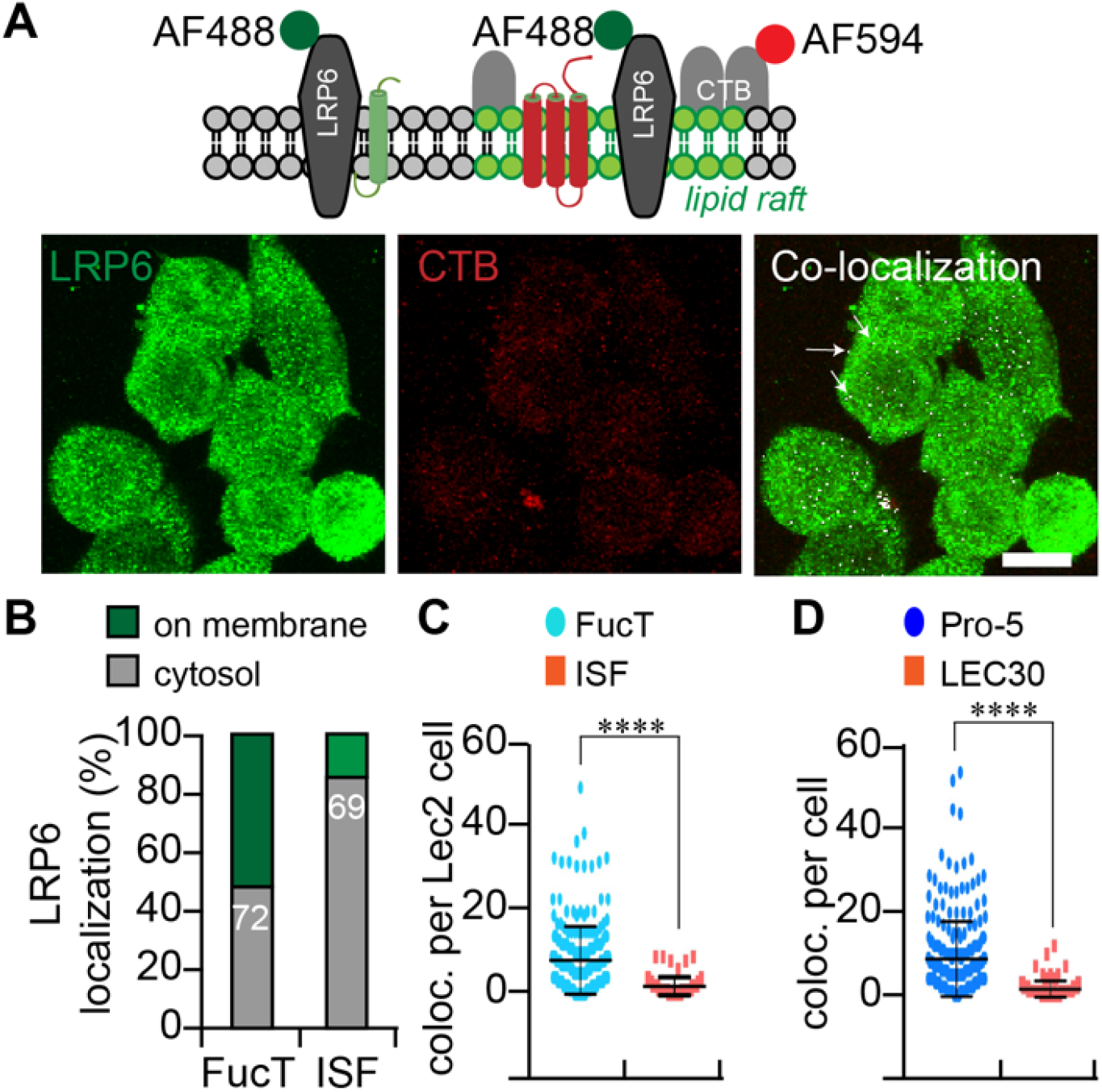
ISF relocalizes LRP6 away from lipid rafts. (A) A schematic representation (top panel) of the membrane topological location of the lipid-raft marker CTB and LRP6. The colocalization of LRP6 and CTB (white dots) in lipid rafts was calculated from confocal images using ImageJ, based on immunofluorescence of antibody to LRP6 (green) and labeled CTB (red). (B) Comparing the colocalization of LRP6 and CTB showed that LRP6 moved from membrane to cytosol in ISF-treated Pro-5 cells. The numbers show the cells counted in this assay. (C, D) Plots of the colocalization of LRP6 and CTB in ISF-treated and FucT-treated Lec2 cells (C) or in untreated LEC30 cells (D). Colocalization was measured by ImageJ and is indicated by white dots. Note: ****, Student t test p<0.0001. Scale bar: 20 μM.

### ISF enhanced the endocytosis of LRP6

To determine if the re-localization of LRP6 was mediated through endocytosis, we examined the co-localization of LRP6 and specific endosomal markers, RAB5, an early endosomal marker; RAB 7, a marker of late endosome (LE),^45^ which is involved in the transfer of cargo to the lysosome for degradation; and RAB 11,^46^ a marker of recycling endosomes (RE). In this experiment, we expressed FLAG-LRP6 in CHO cells and then pulse-labeled the surface LRP6 using anti-FLAG antibody conjugated to Alexa Fluor 488 for 30 min. Then we chased the location of fluorescent-labeled LRP6 after ISF treatment.

We found that ISF increased the colocalization of LRP6 and RAB5-mCherry shortly after the initiation of pulse labeling. However, without ISF-treatment, there was essentially no colocalization of LRP6 and RAB5 (Fig. 6A). By contrast, after ISF we could clearly detect formation of large clusters of colocalized LRP6 and RAB5 (Fig. 6A, lower panel). Subsequently, we analyzed the colocalization of LRP6 with RAB5, RAB7 and RAB 11, respectively, using ImageJ to calculate Pearson’s correlation coefficient (PCC). The value of PCC defines the colocalization between two factors. A PCC value of 1 means total colocalization; 0 means random localization between two locations; and −1 means the two factors are not colocalized. As shown in Fig. 6D, ISF did not induce any notable changes in colocalization between LRP6 and RAB7. However, the colocalization of LRP6 with Rab11 increased significantly by 30 minutes following ISF (Fig. 6E).

**Figure 6.**
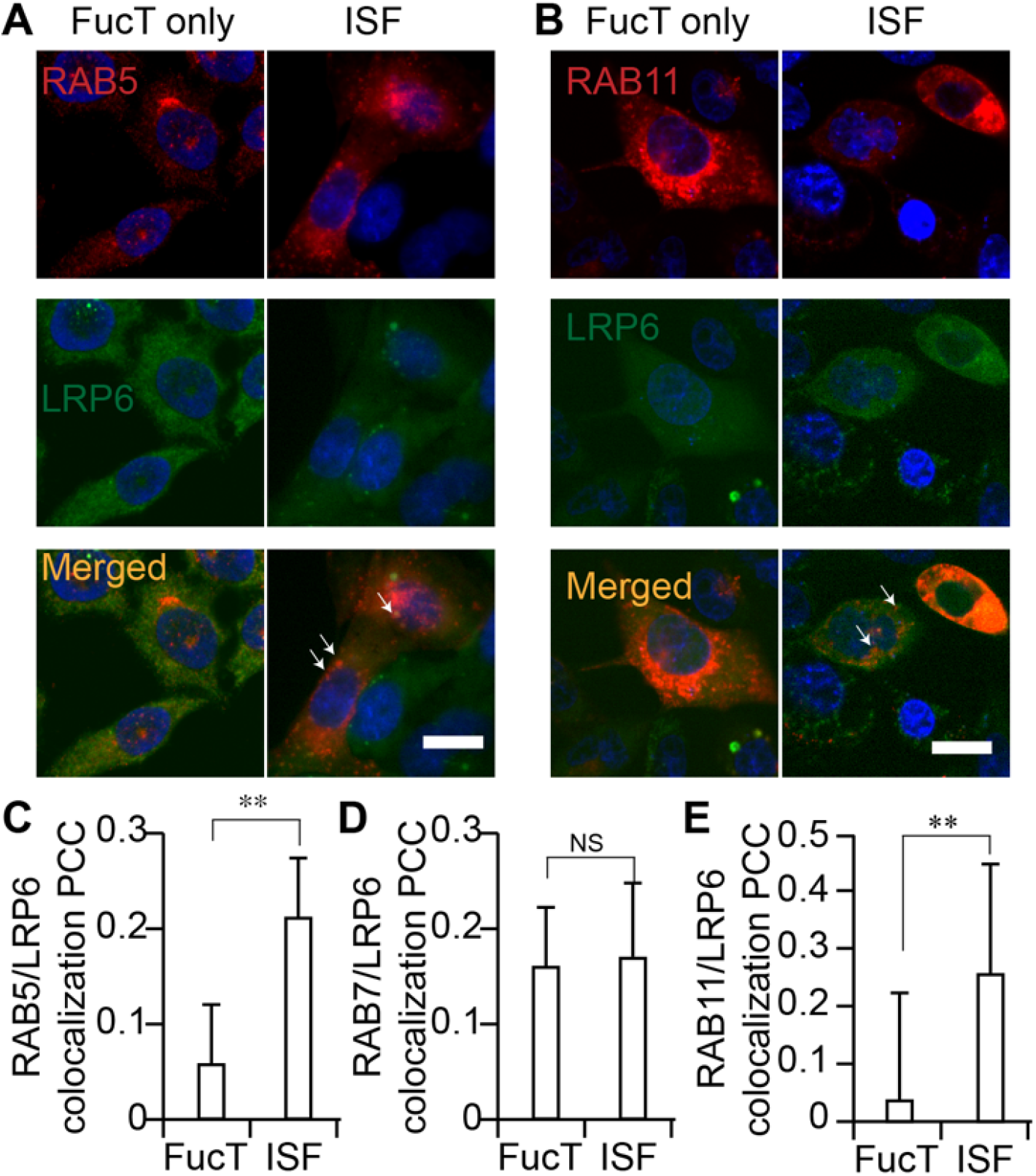
Tracking LRP6 endocytosis induced by ISF. (A, B) Confocal visualization of LRP6 (LRP6-GFP, green) in FucT only-treated and ISF-treated Pro-5 cells. Early endosome marker RAB5-mCherry (A, red) and recycling endosome marker RAB11-mCheery (B, red) were transiently expressed in Pro-5 cells. The colocalization of LRP6 with RAB5 or RAB11 is shown in yellow. Scale bar: 20 μM. (C-E) Colocalization of RAB5 (C), RAB7 (D) or RAB11 (E) with LRP6 in FucT-only-treated and ISF-treated Pro-5 cells, is shown as Pearson’s correlation coefficient (PCC). Note: **, Student’s t test p<0.005; NS, not significant.

### Enhancing endocytosis also increases N-glycan LacNAc fucosylation-mediated Wnt inhibition, whereas decreasing endocytosis rescues the inhibition

To further investigate whether N-glycan LacNAc fucosylation-induced Wnt inhibition occurs through an endocytic process, we sought to artificially enhance endocytosis by over-expressing dynamin-1^36^ or to inhibit this process by either over-expressing dominant-negative (DN) dynamin-1 or using Dynasore, a dynamin-1 inhibitor.^47^ Dynamin-1 is a protein responsible for all vesicle-mediated endocytosis, including clathrin-mediated and caveolin-mediated processes. Under each of these conditions, we quantified the ratio of Wnt activity in ISF-treated Pro-5 cells to that of Pro-5 cells treated with FucT only. Alternatively, under the same condition, we quantified the ratio of Wnt activity in LEC30 to that of Pro-5 cells. This ratio directly reflects the effect of Wnt inhibition. The lower the ratio becomes, the more severe the inhibition of Wnt signaling that occurred.

We observed that transient expression of dynamin-1 decreased the ratio of Wnt activity in the ISF-treated cells to that in Pro-5 cells treated with FucT only, and the decreased amount was positively correlated to the amount of dynamin-1 expressed (Fig. S6A). This result strongly suggested that dynamin-1 expression mimics the difference in Wnt activity between LacNAc N-fucosylated and non-N-fucosylated cells. Accordingly, treatment by Dynasore increased the ratio of Wnt activity from 0.57 to 0.70 (Fig. S6B), a similar result to the Wnt signaling difference decreased between ISF-treated and non-treated cells. Likewise, with DN dynamin-1 expression, the ratio of Wnt activity of LEC30 to that of Pro-5 cells increased from approximately 18% to ~100%, suggesting that near full rescue of N-glycan LacNAc fucosylation-induced inhibition was achieved (Fig. S6C). A similar trend was also observed for the ISF-treated Pro-5 cells (Fig. S6D).

### N-glycan LacNAc fucosylation mediated Wnt inhibition can be rescued by free Fuc

Since increasing cell-surface N-glycan LacNAc fucosylation led to inhibition of Wnt signaling inhibition, we investigated whether adding free Fuc to the culture medium could specifically reverse this inhibition. This experiment was based on the hypothesis that fucosylated LacNAc on ISF-treated cells might be recognized by a Fuc-specific receptor which was binding to ISF-treated LRP6 and facilitating its endocytosis. Free Fuc might compete for this binding, allowing fucosylated LRP6 to remain at the cell surface. Four monosaccharaides, L-Fuc, D-galactose (Gal), D-glucose (Glc) and D-mannose (Man), were used in this experiment. We compared Wnt signaling activity of cells treated with or without ISF in the presence of each free sugars. In Pro-5 (Fig. 7C) and Lec2 (Fig. S7) cells, the addition of free Fuc but not the other monosaccharides rescued ISF-induced inhibition of Wnt signaling. A dose-dependent effect was observed and complete rescue occurred with 750 μM Fuc (Fig. 7D). Likewise, in highly N-fucosylated LEC30 cells, enhanced Wnt signaling activity was observed following the addition of free Fuc but not the other monosacharrides (Fig. S8). Not surprisingly, free Fuc did not block ISF from elevating the cell-surface fucosylation (Fig. S9 top panel) or LRP6 fucosylation as confirmed by Western blot analysis (Fig. S9). Moreover, ISF and Fuc co-treatment did not cause cell-surface LRP6 to decrease (Fig. S10).

**Figure 7.**
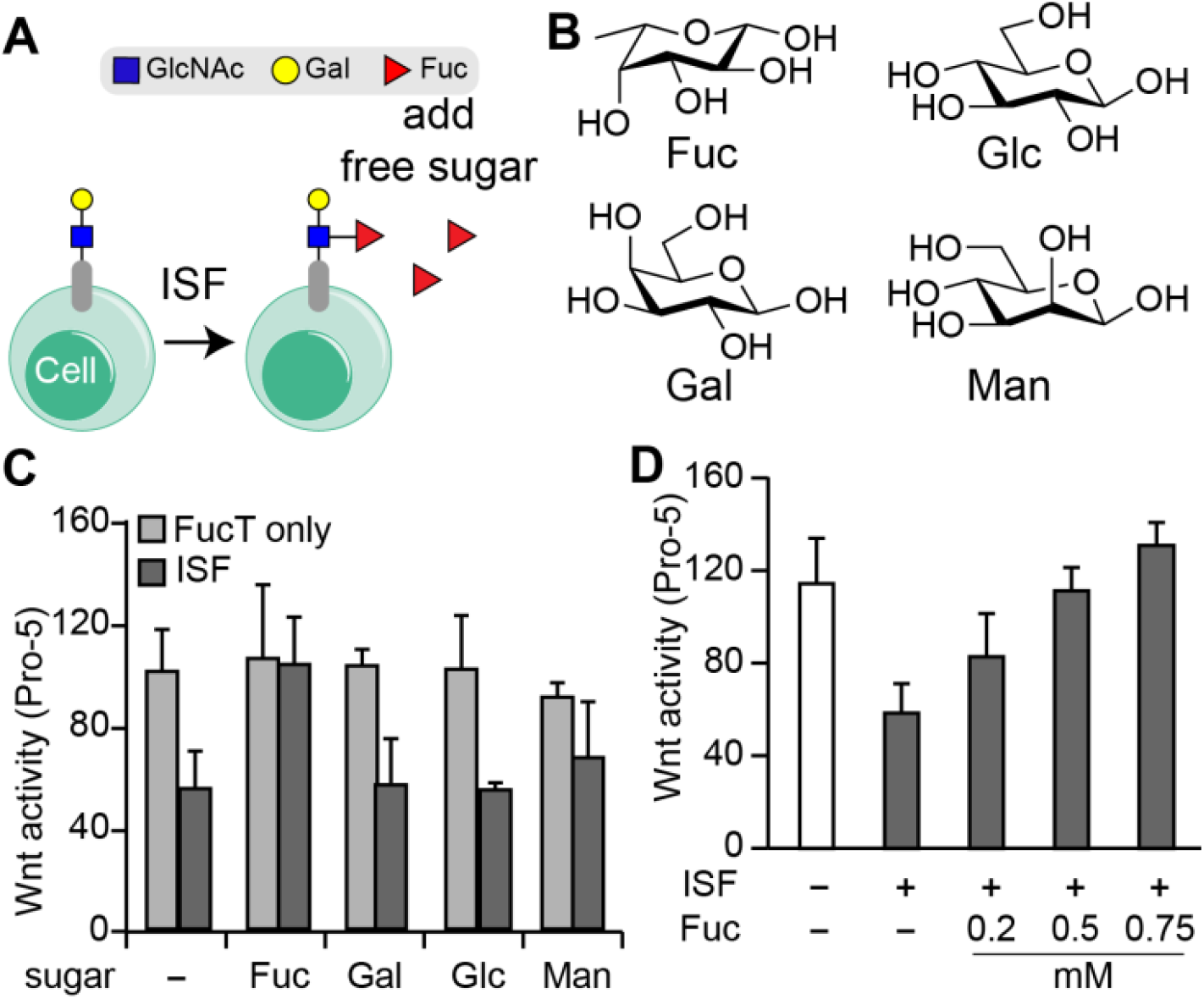
Free Fuc suppresses ISF-induced Wnt inhibition. (A) Schematic flowchart of the monosaccharide treatment. (B) The chemical structure of four monosaccharaides in a vertebrate *N*-glycan, including L-fucose (Fuc), D-glucose (Glc), D-galactose (Gal) and D-mannose (Man). (C, D) Comparison of Wnt activity in Pro-5 cells treated by ISF or FucT only, in the presence or absence of free monosaccharaides. The bars represent the standard error of three biological replicates.

To further explore roles for N-glycan LacNAc fucosylation in endocytosis mediated by fucosylation, we compared the internalization of un-fucosylated LacNAc disaccharide and its α1-3-fucosylated counterpart—Lewis X (Le^X^). We prepared Cy-3 tagged LacNAc and Le^X^ by conjugating Cy3 to the reducing end of LacNAc and Le^X^, respectively. After incubating Pro-5 and LEC30 cells with these Cy3 tagged oligosaccharides, we observed that significantly higher amounts of Cy3-labeled Le^X^ were internalized compared to the Cy3-labeled LacNAc (Fig. S11). This phenomenon became more obvious at higher sugar concentrations. When 50 μM of both sugars were used, 20–40 fold more Le^X^ was internalized. Internalization was more pronounced in Pro-5 cells than in LEC30 cells. In addition, externally added, free Fuc partially decreased Le^X^ internalization, but not internalization of the unfucosylated LacNAc derivative (Fig. S11). When Pro-5 cells were treated with 50 μM of Cy3-Le^X^ together with 1 mM of free Fuc, the internalization of Le^X^ was reduced to half (Fig. S11). Cell-surface ISF treatment also reduced Le^X^-Cy3 internalization (Fig. S12). Together, these results suggest the possible presence of a Fuc binding protein or receptor on the CHO cell surface that is involved in mediating the turnover of fucosylated glycoconjugates (i.e., LRP6) (Fig. S14).

### N-glycan LacNAc fucosylation suppressed Wnt/β-catenin signaling in breast cancer cells

Since Wnt/β-catenin is hyperactivated in triple-negative breast cancer (TNBC), which is believed to promote cancer progression,^48^ we examined whether ISF is applicable to modulate Wnt/ β-catenin signaling in TNBC. To this end, the TNBC MDA-MB-231 cells and non-TNBC MCF-7 breast cancer cells were treated with ISF or not treated. Cellular β-catenin levels were detected by Western blot. As expected, after ISF treatment, we observed a significant decrease of the β-catenin level in both MDA-MB-231 and MCF-7 cancer cells (Fig. 8A). Wound healing assays further revealed that the ISF treatment also impaired the migration of both cancer cells (Fig. 8B and 8C).

**Figure 8.**
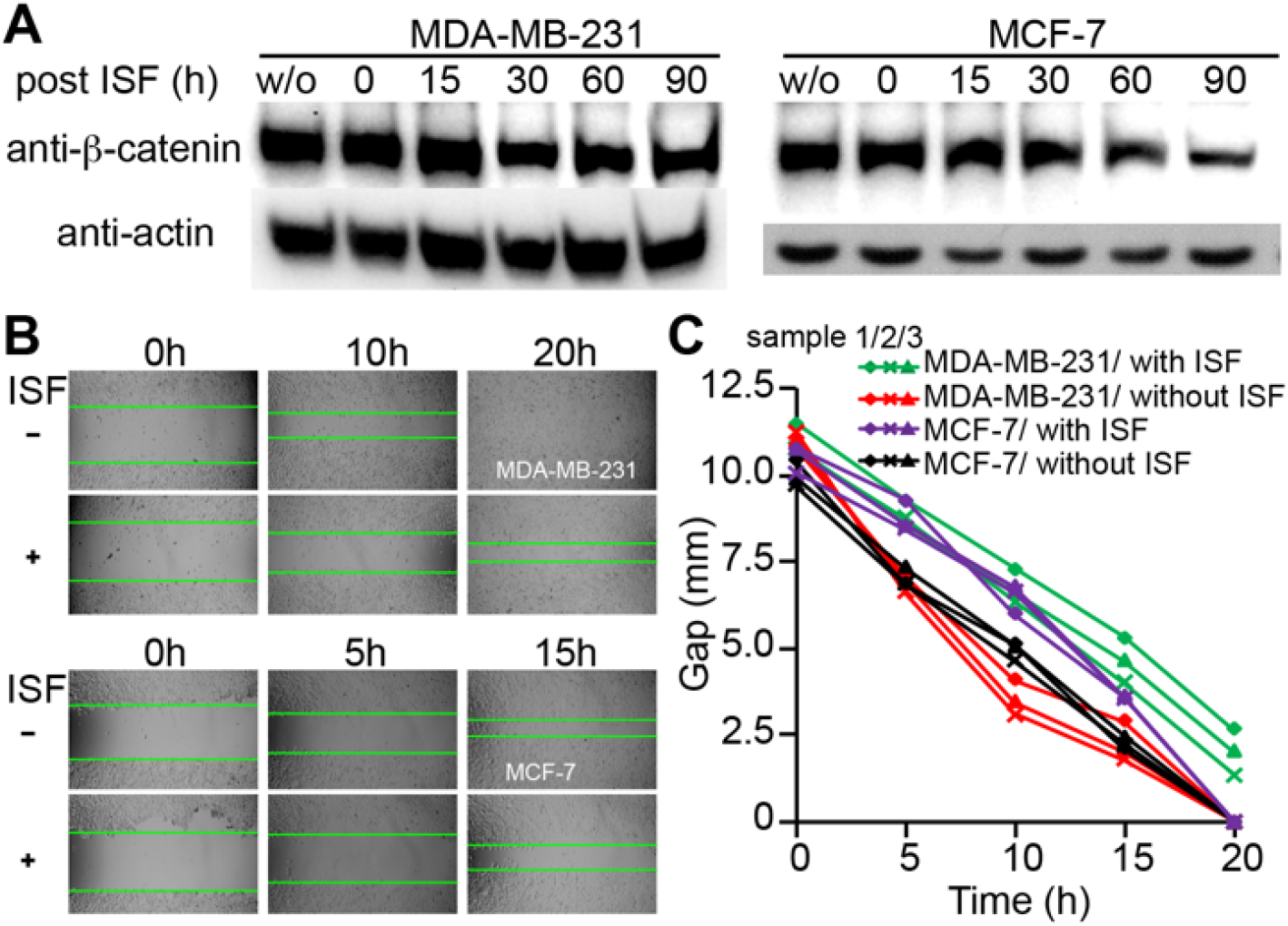
Cell-surface ISF suppresses Wnt/β-catenin signaling in breast cancer cells. (A) Time-dependent decrease of the β-catenin level in ISF-treated MDA-MB-231 and MCF-7 breast cancer cells. Western blot of anti-β-catenin (upper panel) and anti-actin (lower panel). (B) Wound healing assay results from MDA-MB-231 and MCF-7 cells treated with ISF or not. (C) Time-dependent gap close assays of MDA-MB-231 and MCF-7 cells treated with ISF or not. Three biological replicates are presented here.

## Discussion

Here we report the discovery that increasing cell surface N-glycan LacNAc α1-3-fucosylation exacerbates the endocytosis of lipid-raft localized Wnt-receptor LRP6, which in turn suppresses Wnt/β-catenin signaling (Fig. s14). Our results also suggest that the majority of internalized cell surface LRP6 enters the recycling pathway rather than the degradation pathway following endocytosis induced by ISF treatment. Interestingly, the ISF-induced inhibition was completely rescued by externally added free Fuc, suggesting the existence of a fucose receptor involved in the caveolin-mediated endocytosis of LRP6 in membrane lipid rafts. Although further evidence is required to identify this cell-surface Fuc-specific receptor, our discovery suggests that the availability of fucose in the extracellular milieu and N-glycan LacNAc fucosylation mediated by Golgi fucosyltransferases and GDP-Fuc serve as reciprocal switches to modulate Wnt activity. It is well known that LRP6 in complex with WNT-FZD is regulated by phosphorylation.^4,5,40,49^ Upon phosphorylation by several kinases, the cytoplasmic tail of LRP6 recruits AXIN, a scaffold protein that further supports the formation of the LRP-FZD dimer to promote Wnt signaling. The N-glycan LacNAc fucosylation-mediated LRP6 endocytosis we discovered in the current study provides an additional regulatory mechanism for Wnt signaling.

Additionally, Wnt signaling plays a critical role in the etiology of various cancers. For example, aberrant activation of Wnt signaling has also been observed in colorectal cancer^50^ and other gastrointestinal tract cancers^51^. Inhibiting the Wnt pathway has been validated for therapeutic intervention of colorectal cancer^52,53^ as a few anti-cancer drugs that modulate the Wnt pathway in vivo have been approved by the US Food and Drug Administration (FDA)^54^. The results from our study suggest there may be potential for using chemo-enzymatic glycan editing^55^ (i.e., in situ N-glycan LacNAc fucosylation) in cancer therapeutics in order to modulate Wnt activity in a cancer cell. By exploiting targeted drug delivery approaches or *in situ* cancer tissue microinjection, fucosyltransferase and GDP-Fuc could be delivered into the tumor microenvironment to induce cell surface N-glycan LacNAc fucosylation to inhibit Wnt signaling in cancer cells. Such an approach may be useful for treating TNBCs, an aggressive form of cancer that fetches unpredictable and poor prognosis due to limited treatment options.

One outstanding problem is that LacNAc is widely distributed on many membrane glycoproteins. The α1-3-linked Fuc added via ISF may also affect the functions of these proteins^55^. To enhance the specificity of ISF glycan editing toward a particular cell-surface glycoprotein, it will be necessary to develop more precise targeting strategies. Inspired by selective desialylation of the tumor cell glycocalyx guided by an antibody that targets HER2, an antigen highly expressed in tumors,^56^ we envision that the selective N-glycan LacNAc fucosylation of LRP6 in target cancer cells via a fucosyltransferase-antibody conjugate may selectively inhibit Wnt signaling in cancer cells.

## Supporting Information

Additional figures, materials and methods

